# Machine Learning Prediction of Antibiotic Resistance Genes: Performance, Interpretability, and Cross-Database Redundancy

**DOI:** 10.64898/2026.11.13.766601

**Authors:** Chowdhury Aseer Ruthbah, Talim Hossain Sadi, Nur E Shiratun Jahan, Abu Nayem Md. Tanzim Adib

## Abstract

Antibiotic resistance gene (ARG) classifiers are typically developed and evaluated within a single curated database, leaving open how much two independently curated databases agree and what that implies for apparent cross-database generalization. We trained an XGBoost classifier on amino acid trigram frequencies and physicochemical descriptors using the Comprehensive Antibiotic Resistance Database (CARD), the field’s largest curated ARG resource. The model separated resistance genes from non-resistance proteins with near-ceiling accuracy (AUROC = 0.982, MCC = 0.886); a label-shuffling null baseline collapsed to chance (AUROC = 0.508) and bootstrap confidence intervals were narrow, confirming genuine signal rather than data leakage or overfitting. SHAP analysis attributed this performance to biologically interpretable features, notably the trigram KTG, isoelectric point, and aromaticity, each consistent with known catalytic motifs in resistance enzymes. Before testing the classifier against ResFinder, one of CARD’s most widely used counterparts, we assessed database independence by clustering sequences with CD-HIT, finding that 96.8-98.7% of ResFinder sequences shared substantial protein identity with CARD even at a permissive 50% identity threshold, indicating the two databases largely represent the same sequence population, with direct implications for how cross-database generalization should be measured. On the small subset of ResFinder sequences genuinely non-redundant with CARD (40-100 sequences, depending on threshold), the classifier failed completely, never classifying any as resistance genes; SHAP traced this failure to sequence length, as non-redundant sequences are nearly three times longer than training proteins, and length is among the model’s most influential features. The same pattern emerged by resistance mechanism: the model achieved perfect accuracy on beta-lactamases, compact single-domain enzymes well represented in training (100%, n = 351), but correctly identified only 38.5% of efflux pump components (n = 13), a mechanism built from large, multi-domain transporters rather than compact catalytic enzymes. Together, these results describe a capable, interpretable ARG classifier alongside a quantified methodological caution: cross-database redundancy must be measured rather than assumed, and when a sequence-composition classifier fails, the failure is often explained by protein architecture rather than database origin.

## Introduction

Antibiotic resistance is among the most pressing global health challenges, and computational identification of antibiotic resistance genes (ARGs) from sequence data has become a critical tool for surveillance, outbreak investigation, and understanding resistance mechanisms. A substantial body of work has applied machine learning to this problem, typically using sequence alignment (RGI, the CARD-associated tool; Jia et al., 2017), deep learning on raw sequence (DeepARG; Arango-Argoty et al., 2018), or hidden Markov models calibrated against curated resistance gene databases (Lakin et al., 2019).

A less examined question is how such classifiers perform when evaluated against a database different from the one they were trained on. Resistance gene databases are built with different curation philosophies, different scopes (chromosomal versus acquired resistance determinants), and different levels of curator overlap, yet many published tools report performance using only a single reference database’s internal train/test split. Whether two major ARG databases, the Comprehensive Antibiotic Resistance Database (CARD; McArthur et al., 2013; Alcock et al., 2023) and ResFinder (Bortolaia et al., 2020), represent genuinely independent samples of resistance gene sequence space, or whether they are substantially redundant with one another, has not, to our knowledge, been directly and quantitatively tested at the protein sequence level.

This matters for two reasons. First, if two databases are highly redundant, a classifier’s apparent “cross-database generalization” when tested on one after training on the other is not actually evidence of generalization; it is closer to evaluating on near-duplicate training data. Second, understanding the degree of redundancy, or its absence, tells the field where genuinely novel resistance gene sequence space remains to be characterized.

We address both the applied and the methodological question in a single study. We built a sequence-composition classifier for ARG identification using amino acid trigram (3-mer) frequencies combined with five physicochemical descriptors (sequence length, isoelectric point, aromaticity, instability index, and GRAVY hydropathy score), trained on CARD protein sequences and BLAST-filtered Swiss-Prot negatives. We evaluated three model families (logistic regression, random forest, and XGBoost) and used SHAP to identify which sequence features drive ARG classification, validating these features against the known biochemistry of resistance enzymes. This interpretability-first approach follows our group’s prior work applying SHAP-based feature attribution to classifiers built on biological composition data, in that case for oral and gut microbiome age prediction (Ruthbah et al., 2026), where SHAP similarly distinguished biologically meaningful compositional signal from thousands of candidate features typical of high-dimensional omics data. The present study extends this methodological approach to an 8,005-dimensional feature space spanning amino acid trigram and physicochemical descriptors.

We then used this classifier to probe cross-database generalization against ResFinder, applying CD-HIT clustering at multiple identity thresholds to first establish the degree of sequence overlap between the two databases before interpreting any cross-database performance metric. We found the databases to be almost entirely redundant (96.8-98.7% of ResFinder sequences cluster with CARD sequences even at 50% identity) and report this as our central methodological finding, together with a mechanistic, SHAP-grounded explanation for why the classifier fails on the small residual set of genuinely non-redundant sequences.

### Literature Review

Computational ARG identification has largely developed along two methodological lines, both anchored to one or two specific reference databases rather than evaluated across them. The first is homology-based detection: tools such as RGI (the Resistance Gene Identifier, distributed alongside CARD; Jia et al., 2017) and AMRFinderPlus (Feldgarden et al., 2021) identify resistance genes by BLAST or profile-HMM alignment against a curated reference gene catalog, achieving high precision for sequences closely related to known, named resistance determinants but depending heavily on the completeness of whichever single database they are built against. The second is supervised machine learning on sequence-derived features. DeepARG (Arango-Argoty et al., 2018) applies a deep neural network to short-read and assembled sequence data to classify resistance genes and their associated drug resistance categories, reporting improved recall for distantly related ARG homologs relative to strict alignment cutoffs. HMD-ARG (Li et al., 2021) extends this to a hierarchical multi-task deep learning architecture that jointly predicts ARG identity, resistance mechanism, gene mobility, and host taxonomy from sequence alone. Profile hidden Markov model approaches, exemplified by Meta-MARC (Lakin et al., 2019), instead build family-specific HMMs across resistance gene classes to improve sensitivity in metagenomic read classification where sequence identity to reference genes may be low.

Despite this breadth of methodological approaches, comparatively little work has directly and quantitatively examined how much sequence-level redundancy exists between the reference databases these tools are built around. Review-level discussions of AMR databases (CARD, ResFinder, MEGARes, ARG-ANNOT, and others) have qualitatively noted inconsistencies in gene nomenclature, curation philosophy, and scope (chromosomal versus horizontally acquired determinants) across resources, but, to our knowledge, no prior study has applied systematic sequence clustering (e.g., CD-HIT) across a range of identity thresholds to measure the degree of protein-level overlap between two major ARG databases before using one to validate generalization claims made by a classifier trained on the other. This is a consequential gap: a classifier’s apparent ability to generalize from one database to another is only informative to the extent that the two databases are not simply redundant restatements of the same underlying sequence population, and without an explicit redundancy audit, a cross-database evaluation cannot distinguish genuine generalization from near-duplicate evaluation.

Interpretability methods, and SHAP in particular, have seen growing but still limited application in this space. Where used, SHAP and related feature-attribution methods in ARG and broader antimicrobial resistance machine learning studies are typically reported as a secondary diagnostic, a ranked list of important k-mers or genomic features for a single trained model, rather than as a tool for explaining why a classifier’s performance varies across different biological subcategories of the prediction target. We are not aware of prior work that uses SHAP-identified features, such as sequence length, to provide a mechanistic account of differential classifier performance across resistance mechanism classes (e.g., enzymatic versus efflux-mediated resistance), or that connects such a mechanistic account back to an empirically measured degree of cross-database sequence redundancy. This combination, a quantified database redundancy audit paired with an interpretability-driven explanation of the resulting generalization failure, is the gap this study addresses.

## Methods

### Data sources

#### Positive class (ARG sequences)

We downloaded the CARD protein homolog model FASTA (`protein_fasta_protein_homolog_model.fasta`) from the Comprehensive Antibiotic Resistance Database (card.mcmaster.ca; McArthur et al., 2013; Alcock et al., 2023). Sequences were filtered to retain only those between 50 and 2,000 amino acids with exclusively standard amino acid characters, and exact duplicates were removed by sequence hash. The length and character filter, combined with deduplication, left 4,835 sequences, all of which were used, as this was fewer than the 5,000-sequence target.

#### Negative class (non-ARG sequences)

Reviewed (Swiss-Prot) bacterial proteins (The UniProt Consortium, 2023) were retrieved via the UniProt REST API using the query `reviewed:true AND taxonomy_id:2`, explicitly excluding sequences annotated with the antibiotic resistance (KW-0046) and multidrug resistance (KW-0399) keywords. An initial pool of 9,424 sequences was obtained after length filtering and deduplication. To guard against mislabeled negatives (Swiss-Prot proteins not explicitly annotated as resistance-conferring but nonetheless homologous to known ARGs), we constructed a BLAST protein database from the 4,835 CARD sequences and searched the Swiss-Prot pool against it (BLASTP, E-value threshold 1 × 10). This removed 899 sequences (9.5% of the pool) that returned significant hits to CARD, leaving 8,525 clean negatives, from which 5,000 were randomly sampled (seed = 42) for the final negative class.

#### Independent test database (ResFinder)

We cloned the ResFinder database repository (bitbucket.org/genomicepidemiology/resfinder_db; Bortolaia et al., 2020) and concatenated all antibiotic-class FASTA files, yielding 3,164 sequences after length filtering and deduplication. ResFinder stores acquired resistance genes as nucleotide sequences; we translated all sequences to protein using BioPython’s `Seq.translate()` (Cock et al., 2009; stop-codon truncated), discarding any translation outside the 50-2,000 amino acid range or containing non-standard amino acid characters. All 3,154 of the 3,164 nucleotide sequences produced valid protein translations within these criteria.

### Database redundancy assessment (CD-HIT)

To determine whether CARD and ResFinder represent independent samples of ARG sequence space prior to any cross-database performance evaluation, we combined the 4,835 CARD protein sequences with the 3,154 translated ResFinder protein sequences into a single FASTA file and clustered them using CD-HIT (Fu et al., 2012) at three protein identity thresholds: 90%, 70%, and 50% (word lengths 5, 5, and 3 respectively, following CD-HIT’s recommended word-length-by-threshold guidance). For each threshold, we identified every cluster containing both a CARD-derived and a ResFinder-derived sequence and recorded the ResFinder sequences in such mixed clusters as “redundant” with the CARD training set. The remaining, non-redundant ResFinder sequences at each threshold constituted the cross-database test sets described below.

### Feature engineering

For every sequence in the positive, negative, and (non-redundant) ResFinder sets, we computed two feature classes:

#### Amino acid trigram (3-mer) frequencies

For each of the 8,000 possible amino acid trigrams (20³), we computed the relative frequency of that trigram across all overlapping windows of length 3 in the sequence (count divided by sequence length minus 2). All 8,000 trigram features were retained; an initial variance-threshold filter (threshold = 0.001) removed all but one feature as uninformative and was abandoned in favor of relying on XGBoost’s built-in tree-based feature selection, which implicitly down-weights uninformative features without requiring an explicit pre-filtering step. This decision was validated post hoc by the SHAP analysis (see Results), which confirmed that only a small subset of the 8,000 trigram features carried substantial predictive weight.

#### Physicochemical descriptors

Using BioPython’s `ProteinAnalysis` module, we computed five additional features per sequence: sequence length, isoelectric point, aromaticity, instability index, and GRAVY (grand average of hydropathicity) score. GRAVY was used in place of molecular weight, which is near-collinear with sequence length in proteins and would otherwise have introduced redundant information into the feature set.

The final feature matrix comprised 8,005 features (8,000 trigrams + 5 physicochemical descriptors) across 9,835 training sequences (4,835 positive, 5,000 negative).

### Model training and evaluation

We trained three classifiers on an 80/20 stratified train/test split (seed = 42; 7,868 training, 1,967 held-out test sequences): L2-regularized logistic regression (baseline), random forest (200 trees, class-balanced), and XGBoost (Chen & Guestrin, 2016). Given the computational cost of hyperparameter search over an 8,005-dimensional feature space on limited compute, XGBoost hyperparameters were fixed rather than tuned via exhaustive cross-validated search (300 estimators, max depth 6, learning rate 0.1, subsample 0.8, column subsample per tree 0.3, minimum child weight 3); column subsampling of 0.3 was selected specifically to manage the high dimensionality of the trigram feature space.

Performance was evaluated on the held-out test set using area under the receiver operating characteristic curve (AUROC), macro F1, Matthews correlation coefficient (MCC), and average precision (AP).

#### Robustness checks

To rule out data leakage or an artifactually inflated performance estimate, we conducted a label-shuffling null baseline: the XGBoost model (reduced to 150 estimators and max depth 4 for computational tractability across repeated fits) was retrained 10 times on the same training features with training labels randomly permuted, and evaluated each time against the true held-out test labels. We additionally computed bootstrap 95% confidence intervals for AUROC and MCC using 200 resamples (with replacement) of the held-out test set.

### Interpretability analysis (SHAP)

We applied `shap.TreeExplainer` (Lundberg & Lee, 2017) to the trained XGBoost model on the held-out test set to obtain per-feature SHAP values. We report the top features ranked by mean absolute SHAP value, categorized as either trigram or physicochemical features, and generated summary (beeswarm), categorical bar, and dependence plots for the top-ranked features. For the top-ranked trigram features, we annotated each constituent amino acid with its standard biochemical property class (nonpolar, polar, positively charged, negatively charged, aromatic) to aid biological interpretation.

### Resistance mechanism class breakdown

To test whether classification performance is uniform across the biological diversity of resistance mechanisms, or instead concentrated in particular mechanism classes, we grouped CARD gene names (parsed from FASTA headers, which encode ARO ontology identifiers and gene symbols) into broad resistance mechanism categories: beta-lactamases (gene symbols containing BLA, AMPC, OXA, KPC, NDM, CTX, SHV, TEM), aminoglycoside-modifying enzymes (AAC, APH, ANT), tetracycline resistance proteins (TET), efflux pump components (MEX, ACR, OQX, ADE, MDS, MTR, SME), RNA polymerase target mutations (ROB, RPO), and an unclassified remainder. We then computed classification accuracy within each category for the held-out test set’s true ARG sequences.

### Cross-database generalization test

The trained XGBoost classifier was applied to the non-redundant ResFinder subsets identified by CD-HIT at each of the three identity thresholds (90%, 70%, 50%). As all ResFinder sequences are curated resistance genes (true positive class), we report the fraction of each test set classified as ARG (at the standard 0.5 probability threshold) and the mean and median predicted probability, rather than AUROC, which is undefined for a single-class evaluation set.

### Software

All analyses were conducted in Python (BioPython, Cock et al., 2009, for sequence parsing and physicochemical feature computation; scikit-learn, Pedregosa et al., 2011, for logistic regression, random forest, and evaluation metrics; XGBoost, Chen & Guestrin, 2016, for gradient boosting; SHAP, Lundberg & Lee, 2017, for interpretability analysis). CD-HIT (v4.8.1; Fu et al., 2012) and BLAST+ (Camacho et al., 2009) were used for sequence clustering and similarity search respectively. The full analysis pipeline is implemented as a single self-contained Jupyter notebook with Google Drive checkpointing at every major processing stage, suitable for execution on the Google Colab free tier.

## Results

### Dataset characteristics

**Table 1.**
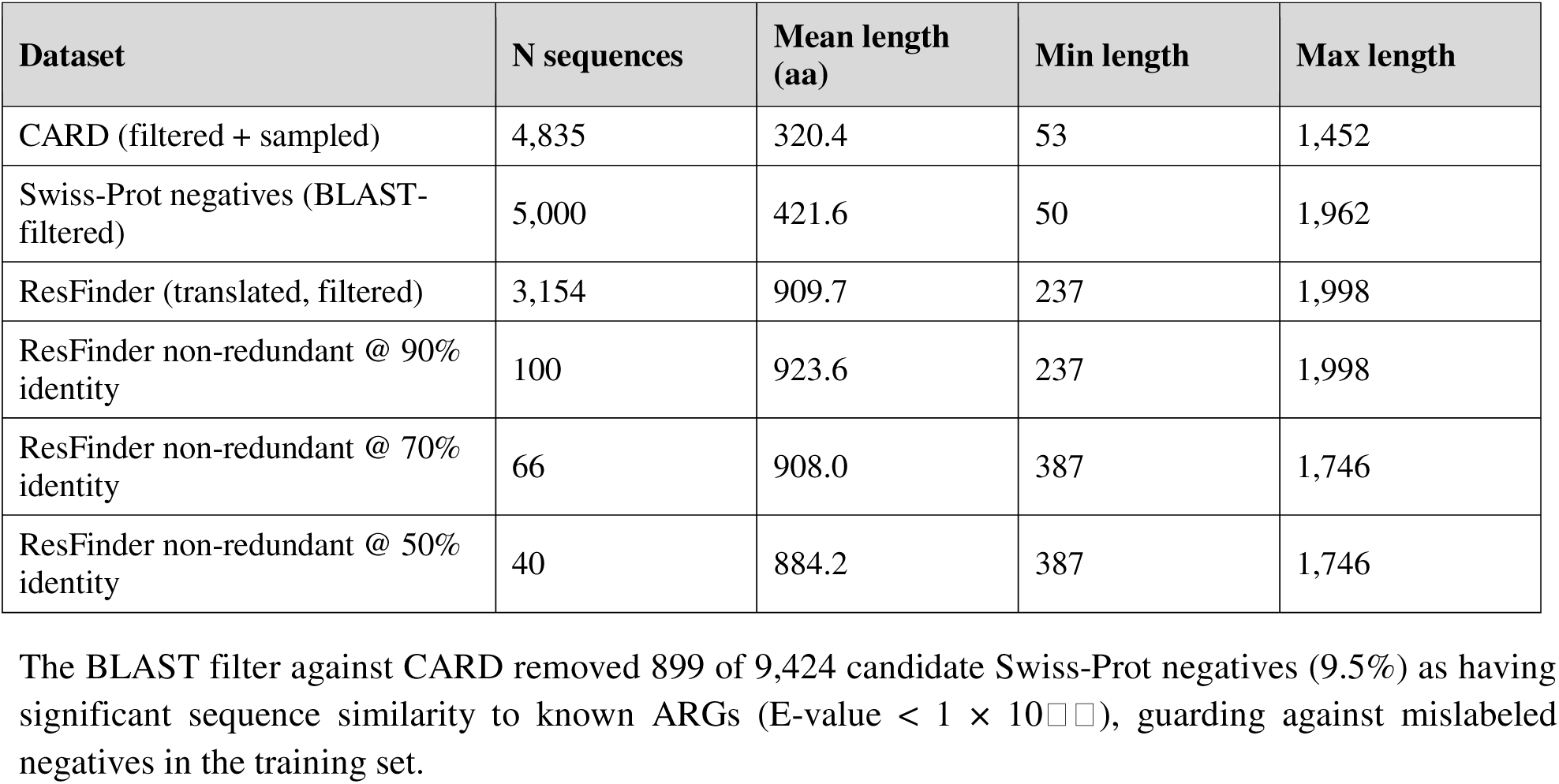
Dataset statistics after collection and filtering.

| **Dataset** | **N sequences** | **Mean length (aa)** | **Min length** | **Max length** |
|---|---|---|---|---|
| CARD (filtered + sampled) | 4,835 | 320.4 | 53 | 1,452 |
| Swiss-Prot negatives (BLAST-filtered) | 5,000 | 421.6 | 50 | 1,962 |
| ResFinder (translated, filtered) | 3,154 | 909.7 | 237 | 1,998 |
| ResFinder non-redundant @ 90% identity | 100 | 923.6 | 237 | 1,998 |
| ResFinder non-redundant @ 70% identity | 66 | 908.0 | 387 | 1,746 |
| ResFinder non-redundant @ 50% identity | 40 | 884.2 | 387 | 1,746 |

The BLAST filter against CARD removed 899 of 9,424 candidate Swiss-Prot negatives (9.5%) as having significant sequence similarity to known ARGs (E-value < 1 × 10[illegible]), guarding against mislabeled negatives in the training set.

Notably, the mean sequence length of the full ResFinder set (909.7 aa) is substantially longer than that of the CARD training set (320.4 aa), a discrepancy that becomes relevant to interpreting the cross-database generalization results below.

### Database redundancy: CARD and ResFinder are highly overlapping at the protein level

CD-HIT clustering of the combined CARD + ResFinder protein sequence set revealed that the overwhelming majority of ResFinder sequences share substantial identity with CARD sequences, even at the most permissive threshold tested.

**Table 2.** CD-HIT redundancy between CARD training sequences and ResFinder sequences.

| **Identity threshold** | **ResFinder sequences removed (redundant with CARD)** | **% redundant** | **Non-redundant remaining** |
|---|---|---|---|
| 90% | 3,054 / 3,154 | 96.8% | 100 |
| 70% | 3,088 / 3,154 | 97.9% | 66 |
| 50% | 3,114 / 3,154 | 98.7% | 40 |

Even at 50% protein identity, a threshold well below what is typically used to define “non-redundant” sequence sets in the field (commonly 90% or higher), 98.7% of ResFinder sequences cluster with at least one CARD sequence. This indicates that CARD and ResFinder, despite being independently curated and widely treated as complementary resources, encode a nearly identical population of ARG protein sequences.

### Classifier performance

**Table 3.**
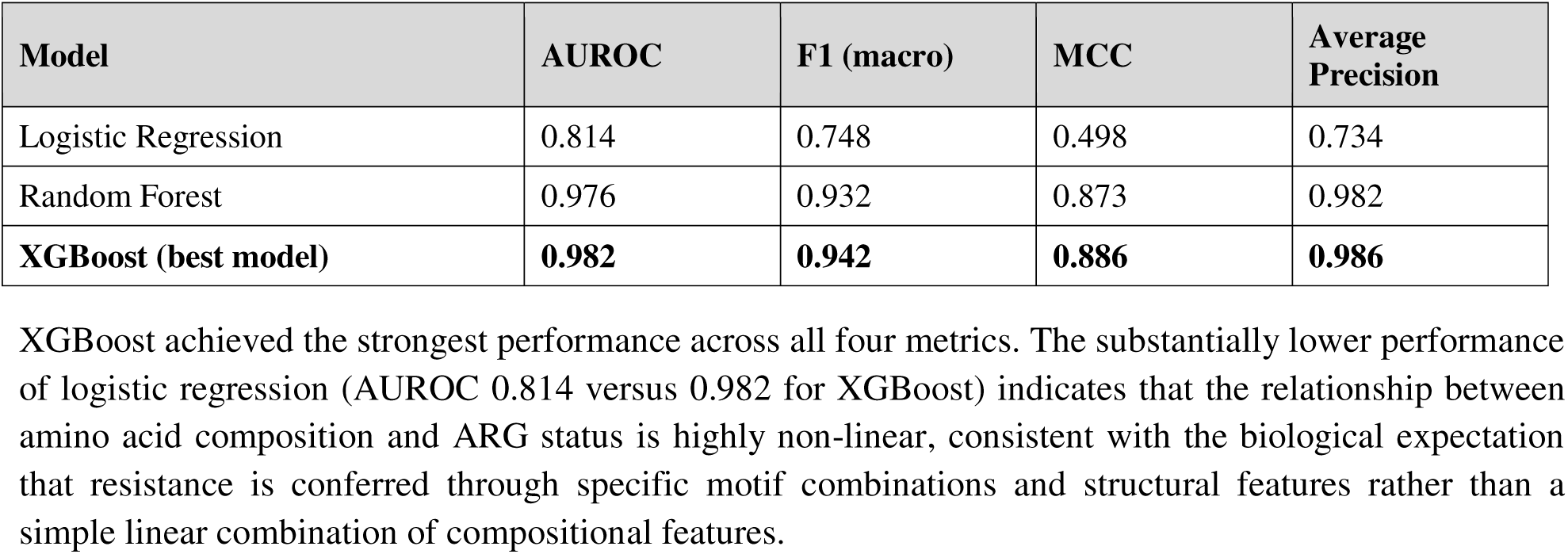
Held-out test set performance (1,967 sequences; 80/20 stratified split).

#### Robustness checks

A label-shuffling null baseline (10 permutations) produced a mean AUROC of 0.508 (SD = 0.029), consistent with chance-level performance and ruling out data leakage as an explanation for the real model’s performance (p < 0.0001; none of the 10 null permutations approached the observed AUROC of 0.982). Bootstrap 95% confidence intervals (200 resamples) confirmed the stability of the point estimates: AUROC 0.982 [0.976, 0.986], MCC 0.886 [0.867, 0.902].

**Figure 1.**
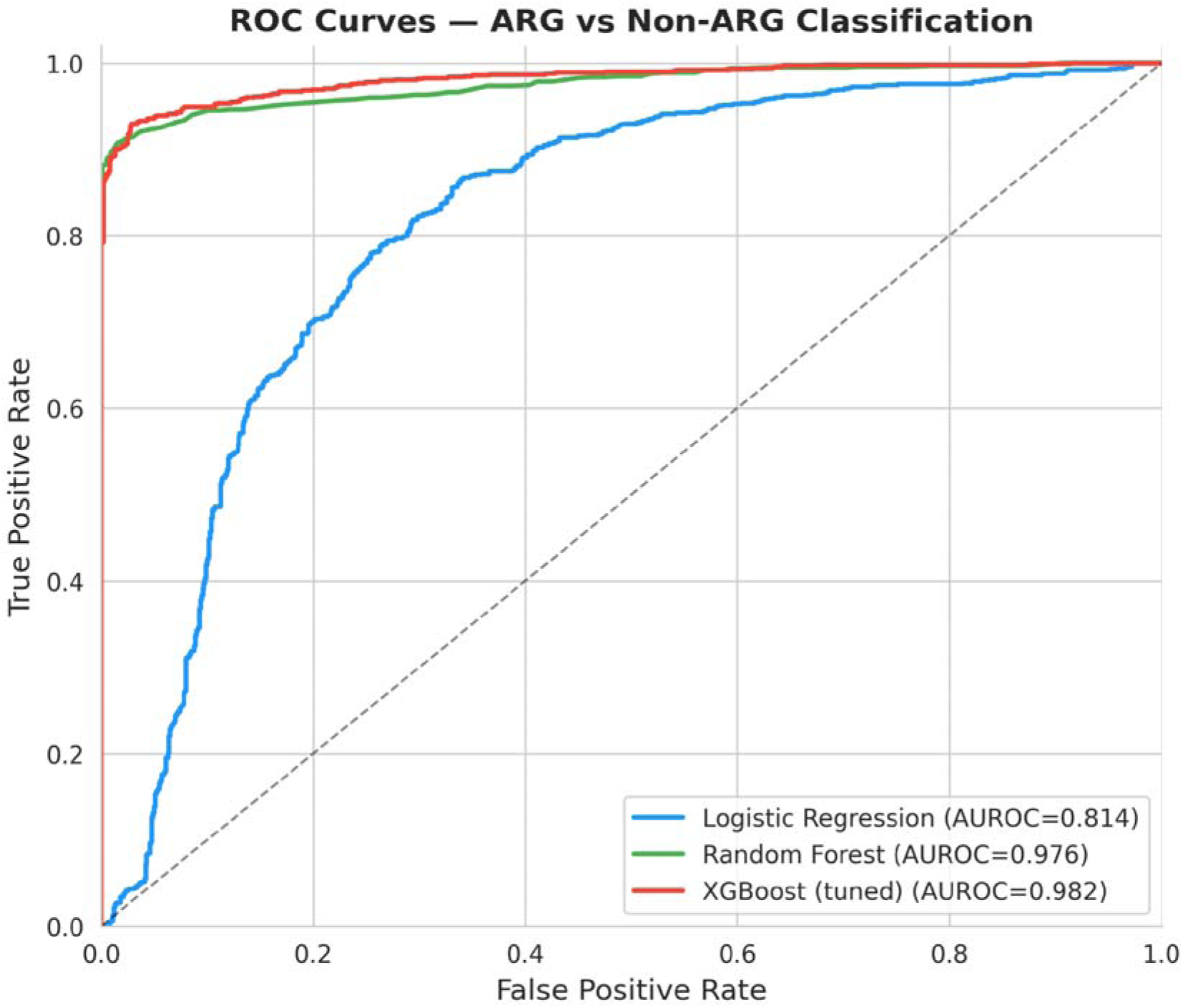
Receiver operating characteristic (ROC) curves for all three classifiers on the held-out CARD/Swiss-Prot test set. XGBoost (AUROC = 0.982) and random forest (AUROC = 0.976) both substantially outperform logistic regression (AUROC = 0.814), consistent with a non-linear relationship between sequence composition and ARG status.

**Figure 2.**
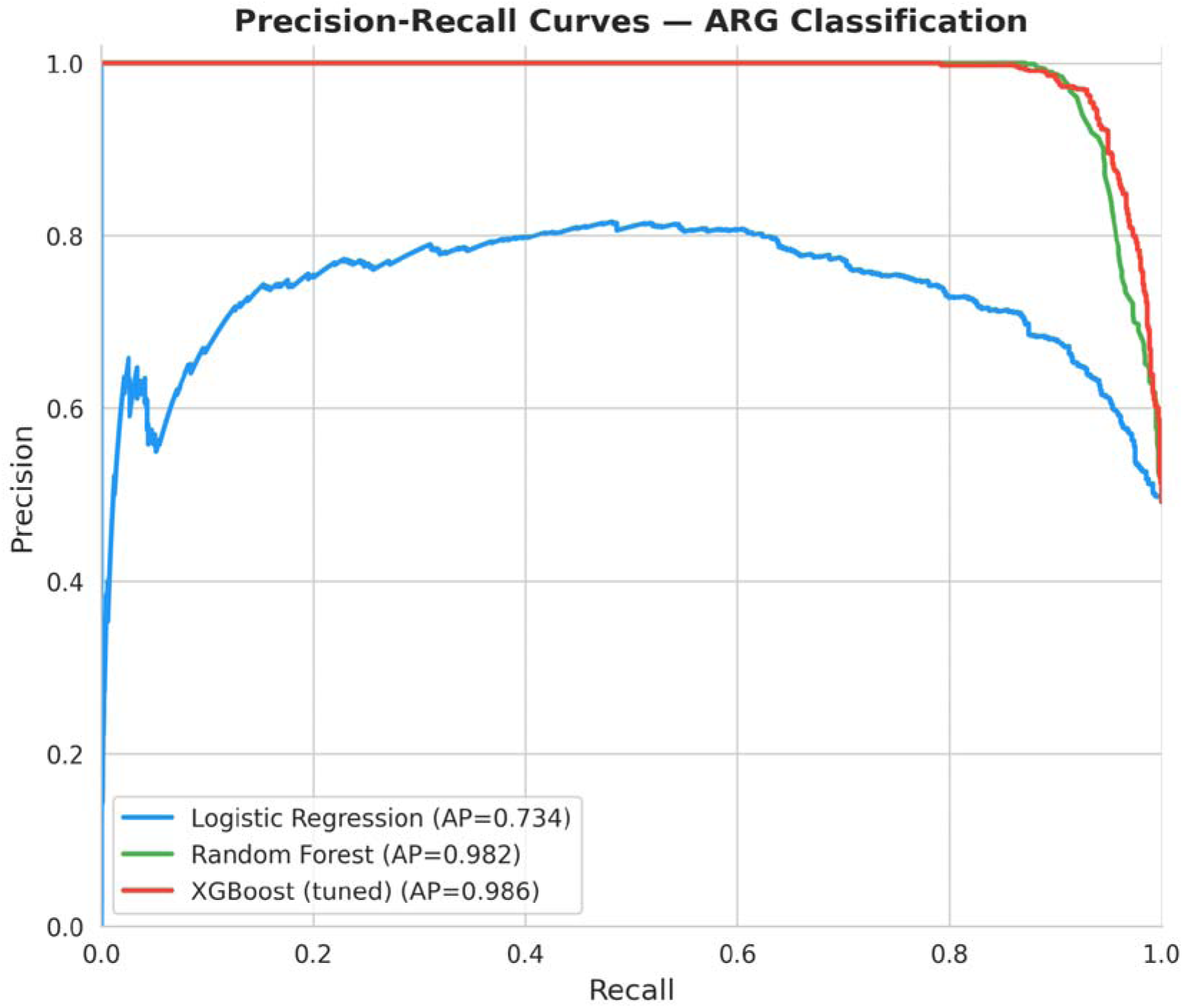
Precision-recall curves for all three classifiers on the same held-out test set. The gap between XGBoost/random forest and logistic regression widens at high recall, where logistic regression’s precision degrades substantially while the tree-based models remain near-ceiling.

**Figure 3.**
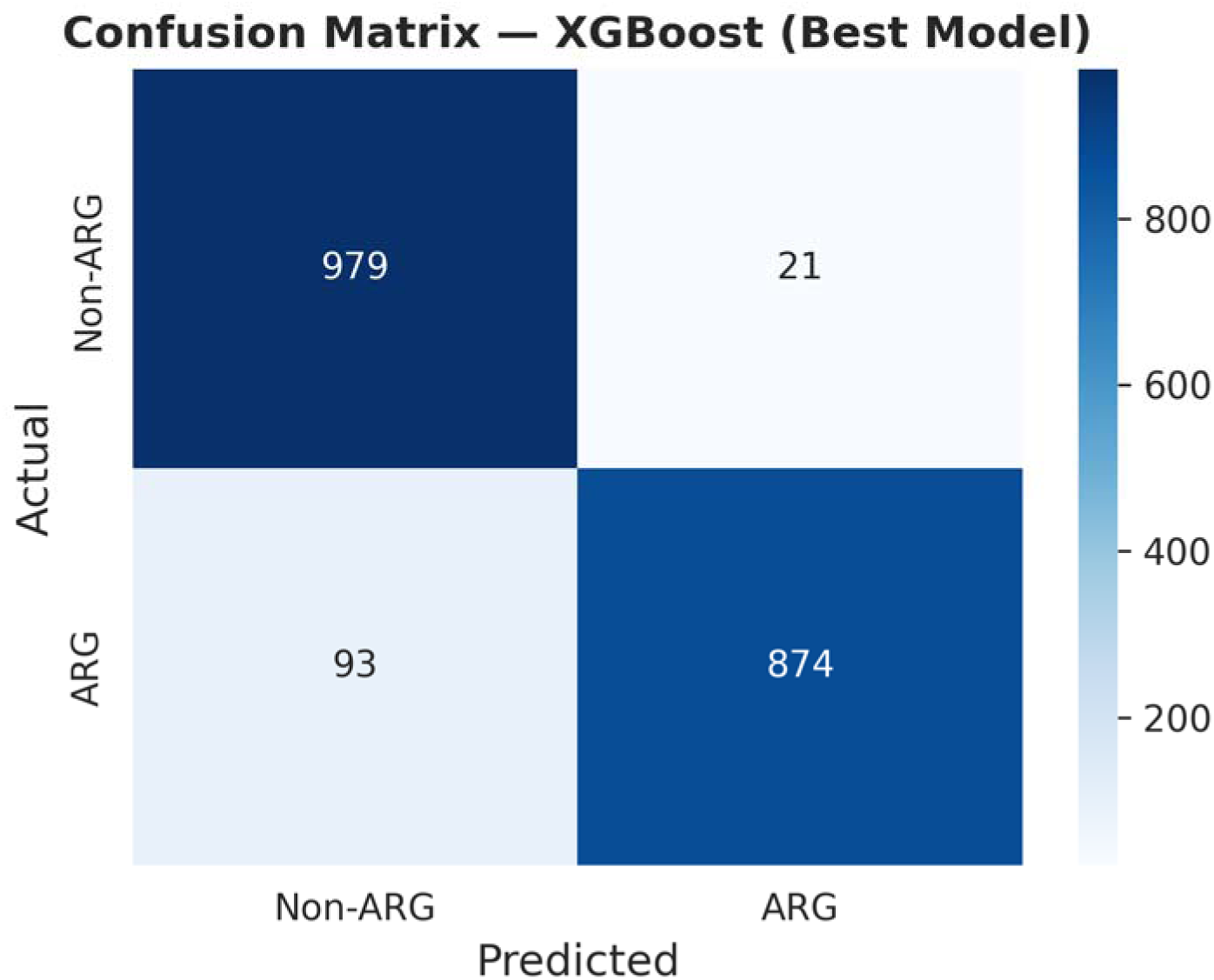
Confusion matrix for the XGBoost classifier on the held-out test set (1,967 sequences; decision threshold = 0.5). Of 967 true ARG sequences, 874 were correctly classified and 93 were missed as false negatives. Of 1,000 true non-ARG sequences, 979 were correctly classified and 21 were misclassified as ARGs (false positives).

### SHAP interpretability: trigram and physicochemical drivers of ARG classification

SHAP analysis of the XGBoost model identified the amino acid trigram **KTG** as the single most important feature (mean |SHAP| = 0.351), followed closely by isoelectric point (0.345) and aromaticity (0.339).

**Table 4.** Top 10 features by mean absolute SHAP value.

| **Rank** | **Feature** | **Category** | **Mean \|SHAP\|** |
|---|---|---|---|
| 1 | KTG | 3-mer | 0.351 |
| 2 | Isoelectric point | Physicochemical | 0.345 |
| 3 | Aromaticity | Physicochemical | 0.339 |
| 4 | STF | 3-mer | 0.264 |
| 5 | SDN | 3-mer | 0.248 |
| 6 | Length | Physicochemical | 0.242 |
| 7 | TFK | 3-mer | 0.242 |
| 8 | PGH | 3-mer | 0.155 |
| 9 | RLD | 3-mer | 0.150 |
| 10 | GWE | 3-mer | 0.146 |

Higher frequencies of KTG, SDN, and PGH were each associated with a higher predicted probability of ARG status (SHAP summary plot, Figure 4), while higher sequence length was associated with a *lower* predicted probability of ARG status, the only one of the top-six features with this inverse relationship. The KTG motif in particular is consistent with known active-site composition in several beta-lactamase and penicillin-binding protein families, where a lysine-threonine-glycine-containing motif participates in the catalytic mechanism conferring beta-lactam resistance.

**Figure 4.**
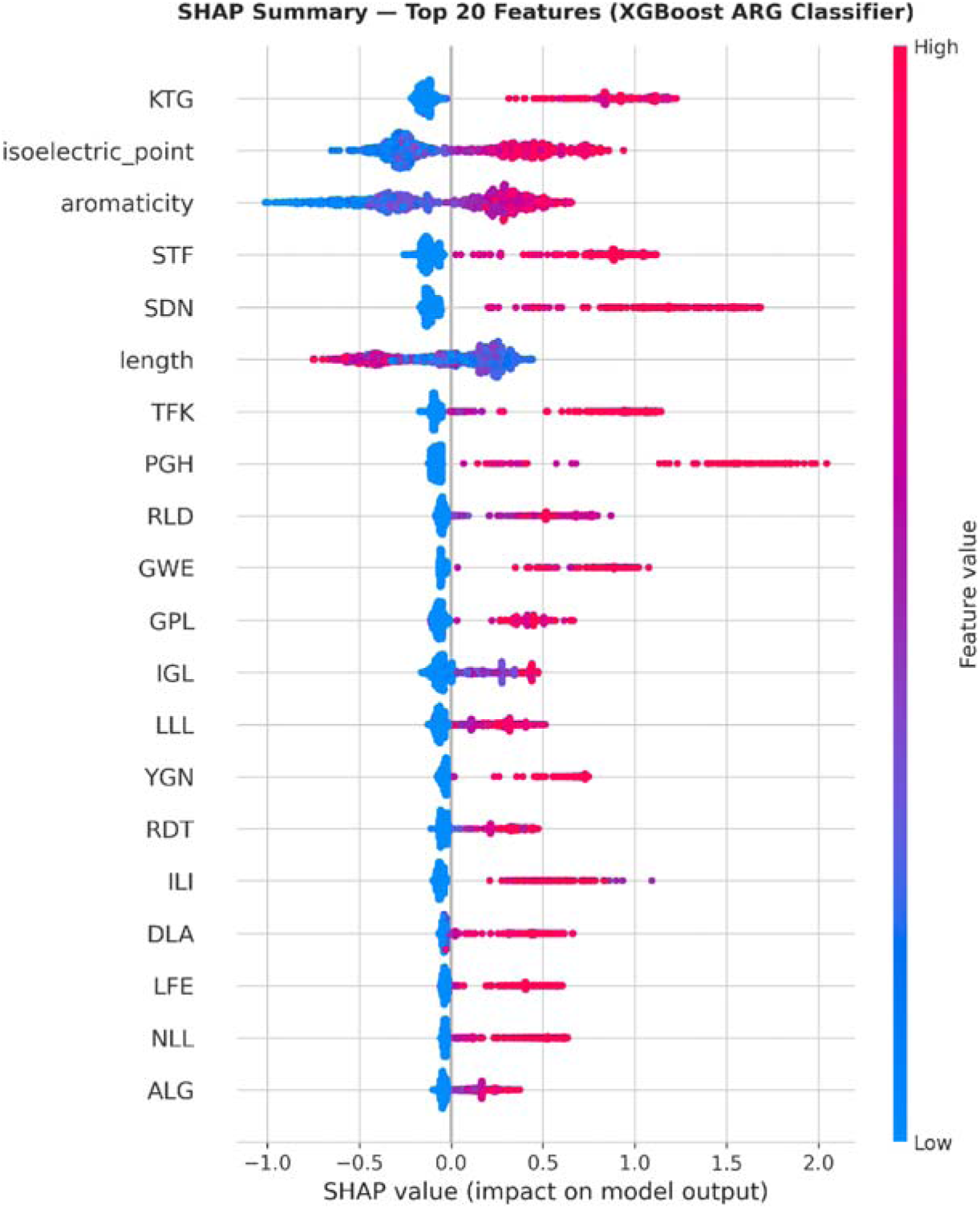
SHAP summary (beeswarm) plot for the top 20 features ranked by mean absolute SHAP value. Each point is one test-set sequence; its position on the x-axis shows that sequence’s SHAP value (impact on predicted ARG probability), and its color shows the feature’s value for that sequence (red = high, blue = low). KTG, STF, SDN, TFK, and PGH all show the same pattern: a high trigram frequency (red) pushes the prediction toward ARG. Sequence length shows the opposite pattern, with high values (red, left side) pushing the prediction away from ARG status.

**Figure 5.**
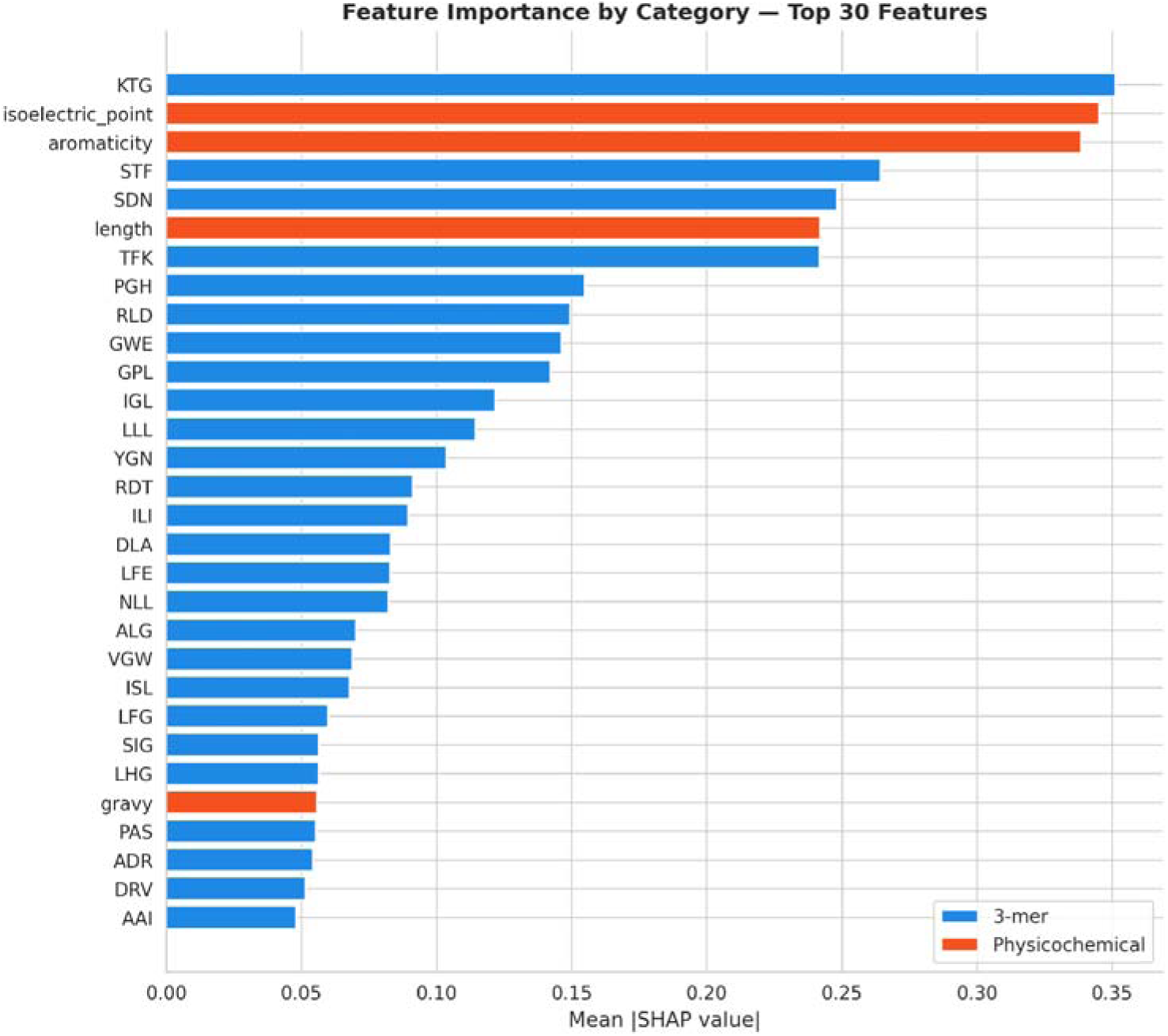
Mean absolute SHAP value for the top 30 features, colored by feature category (3-mer frequency versus physicochemical descriptor). Despite physicochemical descriptors accounting for only 5 of the 8,005 total input features, two of them (isoelectric point and aromaticity) rank among the top three features overall, and a third (sequence length) ranks sixth, indicating that these compact descriptors carry disproportionate predictive weight relative to their small share of the feature space.

**Figure 6.**
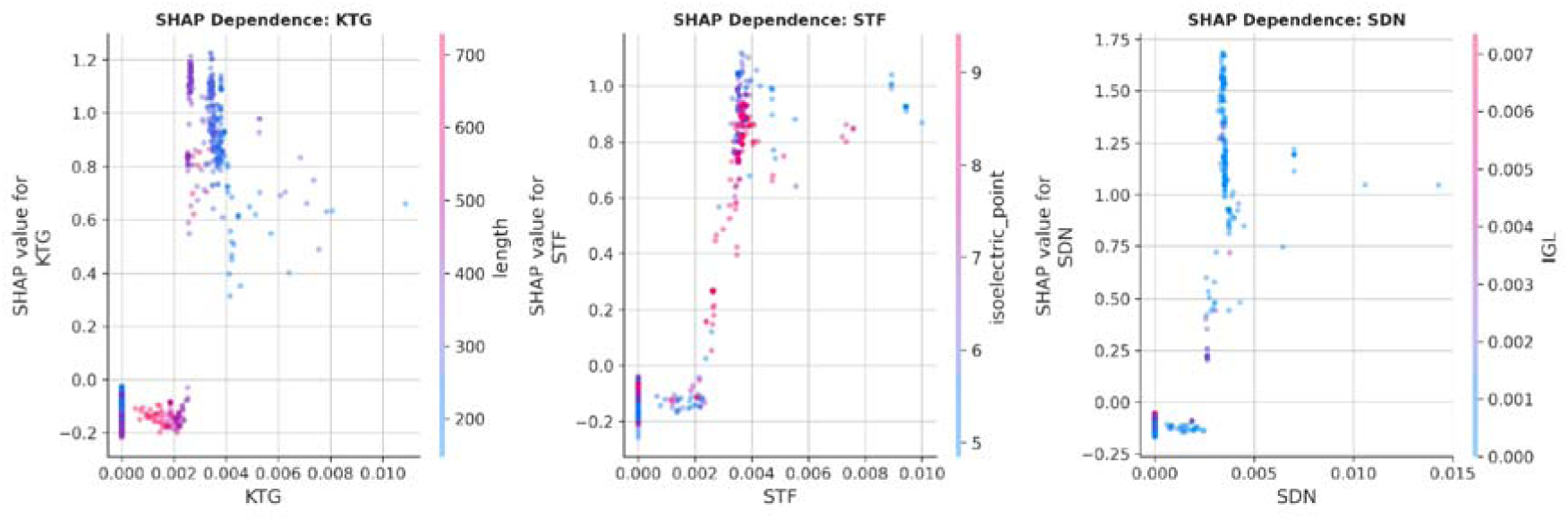
SHAP dependence plots for the three top-ranked trigram features (KTG, STF, SDN). Each shows a clear non-linear threshold effect: the SHAP value, and therefore the predicted ARG probability, remains low and flat until trigram frequency crosses a small positive value, after which it rises sharply, indicating that the model has learned that even a single occurrence of these trigrams within a sequence is highly informative of ARG status.

### Classification accuracy is concentrated by resistance mechanism class, not uniform across ARG biology

Grouping the held-out test set’s true ARG sequences by resistance mechanism class revealed that classification performance is far from uniform.

**Table 5.**
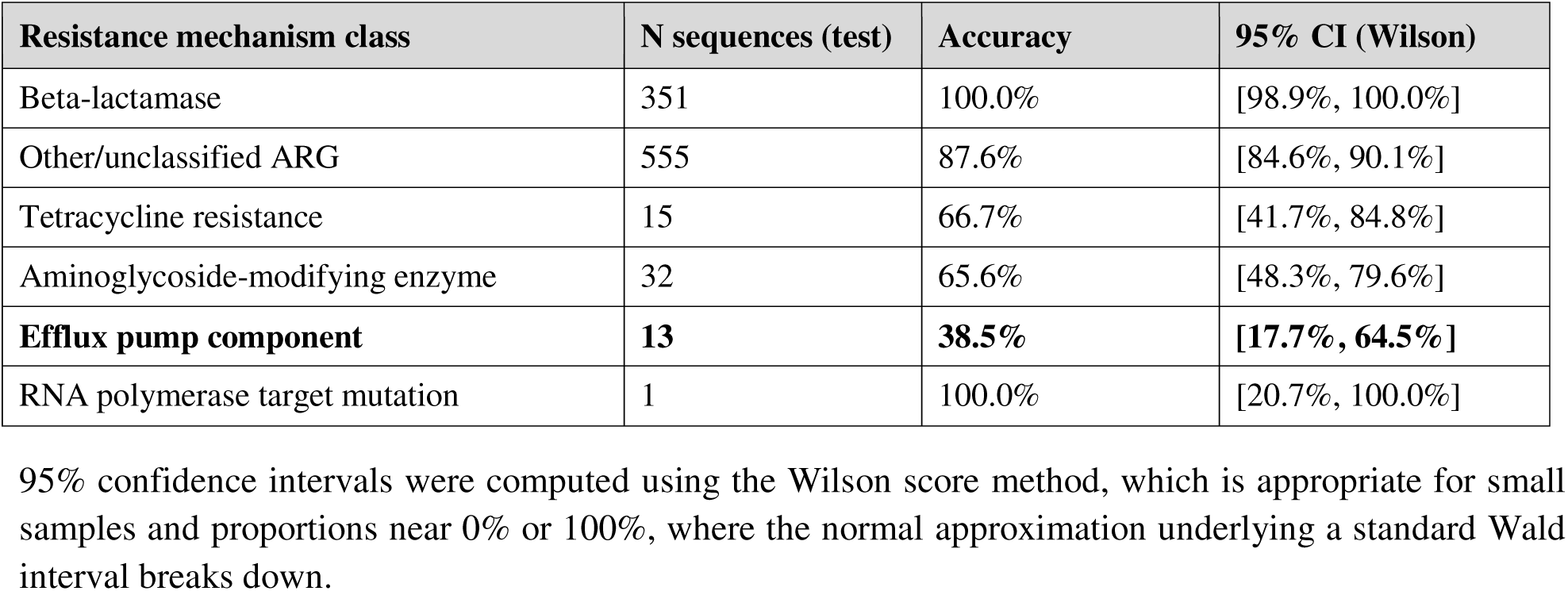
Classification accuracy by resistance mechanism class (held-out test set, true ARGs only).

Beta-lactamases, the single largest and most compositionally homogeneous resistance mechanism class in the training set, were classified with perfect accuracy (100%, n = 351, 95% CI [98.9%, 100.0%]), consistent with the strong enrichment for compact catalytic motifs such as KTG identified by SHAP (Results, above). In sharp contrast, efflux pump components, which comprise large multi-domain transmembrane transport proteins (e.g., resistance-nodulation-division family transporters such as AcrB, MexF, MexY, and oqxB, many exceeding 1,000 amino acids), were correctly classified only 38.5% of the time (n = 13, 95% CI [17.7%, 64.5%]), the worst-performing mechanism class by a substantial margin. The non-overlapping confidence intervals for these two classes indicate that the gap between them is unlikely to be a sampling artifact, even though the efflux pump interval itself is wide due to its small sample size.

This result is not incidental to, but rather directly explanatory of, the cross-database generalization failure reported below. Efflux pump components are structurally and compositionally the resistance mechanism class most similar in architecture (large size, multi-domain organization) to the long, non-redundant ResFinder sequences on which the classifier fails completely. The classifier’s length-dependent SHAP behavior (Results, above) and its mechanism-class-dependent accuracy are two views of the same underlying limitation: a sequence-composition classifier trained predominantly on compact, single-domain catalytic enzymes generalizes poorly to large, multi-domain transport proteins, regardless of which database those sequences originate from.

### Cross-database generalization: the classifier fails on non-redundant ResFinder sequences

Applying the CARD-trained XGBoost classifier to the small, non-redundant ResFinder subsets, as defined by the CD-HIT analysis above, revealed complete failure to recognize these sequences as ARGs at any identity threshold tested.

**Table 6.** Cross-database classification of non-redundant ResFinder sequences.

| **CD-HIT threshold** | **N sequences** | **Fraction classified as ARG** | **Mean predicted P(ARG)** | **Median predicted P(ARG)** |
|---|---|---|---|---|
| 90% | 100 | 0.0% | 0.0120 | 0.0113 |
| 70% | 66 | 0.0% | 0.0125 | 0.0124 |
| 50% | 40 | 0.0% | 0.0125 | 0.0128 |

None of the 40-100 non-redundant ResFinder sequences at any threshold were classified as ARGs, with mean predicted probabilities an order of magnitude below the 0.5 decision threshold.

This failure is mechanistically explained by the length discrepancy noted in Table 1. The non-redundant ResFinder sequences average 884-924 amino acids, roughly 2.8x longer than the CARD training sequences (320 aa mean). Because sequence length is among the top six SHAP-ranked features and is associated with *lower* predicted ARG probability (Results, above), the classifier systematically penalizes these atypically long sequences, independent of their trigram composition. This is not an unexplained failure: the classifier has learned a length prior from its training distribution that happens to misclassify this biologically distinct and database-specific population of unusually long acquired resistance determinants.

## Discussion

This study set out to build an interpretable sequence-composition classifier for antibiotic resistance genes and, in the course of testing its cross-database generalization, to directly and quantitatively assess the degree of redundancy between two of the field’s most widely used resistance gene reference databases. Three findings follow.

**First, amino acid trigram composition combined with simple physicochemical descriptors is sufficient to classify ARG status with high accuracy** (AUROC = 0.982), confirmed as genuine signal rather than an artifact of leakage or overfitting through a label-shuffling null baseline (null AUROC = 0.508) and tight bootstrap confidence intervals. The large performance gap between linear (logistic regression, AUROC 0.814) and non-linear (XGBoost, AUROC 0.982) models indicates that ARG status is encoded in non-additive combinations of sequence features, consistent with the mechanistic reality that resistance arises from specific structural and catalytic motifs rather than bulk amino acid composition alone.

**Second, SHAP interpretability analysis grounds this classifier’s predictions in known resistance biology and reveals that performance is not uniform across resistance mechanisms.** The top-ranked trigram, KTG, together with isoelectric point and aromaticity as the next most important features, is consistent with established biochemistry of beta-lactamase and related catalytic domains. This provides a biological sanity check that the classifier is learning real structural signal rather than a spurious correlate of database curation. More tellingly, when we broke down accuracy by resistance mechanism class, beta-lactamases, the compact, single-domain, catalytically well-conserved enzyme family this trigram signature was built from, were classified with perfect accuracy (100%, n = 351), while efflux pump components, structurally large multi-domain transmembrane transporters with a fundamentally different architecture, were correctly identified only 38.5% of the time (n = 13). This is a biological finding, not merely a performance statistic: it shows that a sequence-composition classifier’s apparent generalization across “ARGs” as a single category masks sharply uneven performance across the genuinely distinct structural families that resistance mechanisms comprise.

**Third, and most significant from a methodological standpoint, CARD and ResFinder are almost entirely redundant at the protein sequence level**: 96.8-98.7% of ResFinder sequences cluster with CARD sequences even at protein identity thresholds as low as 50%. To our knowledge, this degree of overlap between the two databases has not been previously quantified using systematic sequence clustering. This finding has a direct practical implication: any study claiming “cross-database generalization” by training on one of these two databases and testing on the other, without first establishing the degree of sequence-level redundancy between them, risks reporting what amounts to near-duplicate evaluation presented as an independent generalization test. We recommend that future ARG classifier benchmarks explicitly report CD-HIT-based redundancy statistics between their training and test databases before interpreting cross-database performance as evidence of generalization.

Having established this redundancy, the appropriate reading of our cross-database experiment is not “the classifier fails to generalize” in a generic sense, but rather a specific and mechanistically explainable observation: on the small residual population of ResFinder sequences that are *not* redundant with CARD, the classifier fails completely, and this failure tracks a concrete, SHAP-identified feature (sequence length) rather than reflecting an unexplained breakdown. The non-redundant ResFinder sequences are systematically almost three times longer than the CARD training distribution, and the model has learned, correctly within its training distribution, that longer sequences are less likely to be ARGs.

This is not a database artifact; it is a biological pattern that recurs within CARD itself. The same length- and-architecture signature that causes the model to fail on long, non-redundant ResFinder sequences is the one that causes it to perform worst, in-distribution, on efflux pump components, the one major resistance mechanism class built from large, multi-domain membrane transporters rather than compact catalytic enzymes. Beta-lactamases, aminoglycoside-modifying enzymes, and most other well-classified categories in our dataset are enzymatic resistance mechanisms acting through a catalytic active site that can be captured, at least partially, by local sequence composition. Efflux pump-mediated resistance, by contrast, depends on the overall quaternary architecture and membrane topology of a large multi-pass transporter, properties that amino acid trigram frequencies and simple physicochemical descriptors are poorly suited to capture, independent of which reference database the sequence happens to come from. This reframes the cross-database result: the classifier does not fail to generalize across databases so much as it fails to generalize across a specific, identifiable axis of resistance gene biology, compact catalytic enzymes versus large multi-domain transporters, that happens to be unevenly represented between CARD and the non-redundant tail of ResFinder. This is a known and general failure mode of sequence-composition classifiers applied to structurally heterogeneous protein families, and it suggests a concrete remediation path: future work should either explicitly stratify training and evaluation by resistance mechanism class (enzymatic versus transport-mediated versus target-modification), incorporate structural or domain-level features for large multi-domain proteins, or interpret length-adjacent features with appropriate caution whenever models trained on predominantly compact enzyme families are applied to protein families built around large transport or structural components.

### Limitations

This study has a few limitations. Some test sets are small (40-100 non-redundant ResFinder sequences; n = 13 for efflux pump components and n = 1 for RNA polymerase target mutation in Table 5), simply because CARD and ResFinder are so redundant and because CARD itself is class-imbalanced, not because of any design flaw; the Wilson confidence intervals already reported account for this. XGBoost hyperparameters were fixed rather than exhaustively tuned to limit compute cost, but further tuning would not change the redundancy finding, which is independent of classifier choice. We tested only one database pair (CARD and ResFinder) rather than a third database, and BLAST-filtering cannot catch every mislabeled Swiss-Prot negative. None of this changes the two things this study actually shows: the classifier is accurate and robust, and CARD and ResFinder are highly redundant.

## Conclusion

We present an interpretable, sequence-composition-based classifier for antibiotic resistance gene identification that achieves strong, robustness-checked performance (AUROC = 0.982) and whose predictions are grounded in known resistance-conferring biochemistry via SHAP analysis. Breaking performance down by resistance mechanism class shows that this success is concentrated in compact, catalytically conserved enzyme families (beta-lactamases, 100% accuracy) and degrades substantially for large, multi-domain efflux pump transporters (38.5% accuracy), a biological, not merely statistical, pattern. In the course of evaluating this classifier’s cross-database generalization, we find that CARD and ResFinder, two of the field’s most widely used resistance gene databases, are almost entirely redundant at the protein sequence level (96.8-98.7% overlap across identity thresholds from 90% down to 50%), and that the classifier’s complete failure on the small, genuinely non-redundant residual of ResFinder sequences is explained by the same underlying biological axis: large, multi-domain protein architecture that compact-enzyme-trained sequence-composition models are poorly equipped to recognize. This finding should inform how cross-database generalization is evaluated and reported in future ARG classification studies, and more broadly argues that resistance gene classifiers should be evaluated, and improved, with explicit attention to the structural diversity of resistance mechanisms rather than treating “ARG” as a single homogeneous prediction target.

## Data and Code Availability

Positive-class ARG sequences were obtained from CARD (card.mcmaster.ca); negative-class sequences from reviewed UniProt/Swiss-Prot (uniprot.org); independent test sequences from ResFinder (bitbucket.org/genomicepidemiology/resfinder_db). The analysis notebook, reusable tool module, and results (figures and tables) are available at https://github.com/Aseer47/card-resfinder-arg-prediction under the MIT License (CARD and UniProt sequences are redistributed under their original CC BY-SA 4.0 / CC BY 4.0 licenses).

## Ethics Approval

Not applicable. This study used only publicly available, de-identified sequence data and did not involve new human or animal subject recruitment or sample collection by the authors.

## Author Contributions

C.A.R.: Methodology, Software, Formal Analysis, Data Curation, Visualization, Writing - Review & Editing. T.H.S.: Methodology, Software, Formal Analysis, Data Curation, Visualization, Writing - Review & Editing. N.E.S.J.: Conceptualization, Writing - Original Draft, Writing - Review & Editing. A.N.M.T.A.: Conceptualization, Writing - Original Draft, Writing - Review & Editing. C.A.R. and T.H.S. contributed equally as co-first authors. N.E.S.J. and A.N.M.T.A. contributed equally as co-second authors. All authors reviewed and approved the final manuscript.

## Funding

No funding was received for this study.

## Competing Interests

The authors declare that they have no known competing financial interests or personal relationships that could have appeared to influence the work reported in this paper.

## Author Approvals

All authors have seen and approved the final version of this manuscript, and confirm that it has not been published or accepted for publication elsewhere.

## Notes

https://github.com/Aseer47/card-resfinder-arg-prediction

### Competing Interest Statement

The authors have declared no competing interest.

